# LSm4 methylation contributes to hydrogen sulfide-mediated stomatal closure

**DOI:** 10.64898/2026.11.14.766680

**Authors:** SB Torres, D Scuffi, R Pantaleno, YC Agrofoglio, MB De Luca, D Staiger, C Garcia-Mata, JL Mateos, MJ Iglesias

## Abstract

Post-translational modifications (PTMs) expand proteomic diversity, yet how redox PTMs intersect with post-transcriptional regulation remains poorly understood. Hydrogen sulfide (H_2_S) signals partly through cysteine persulfidation. Here, we investigated the relationship among persulfidation, protein arginine methyltransferase 5 (PRMT5)-mediated methylation, and alternative splicing in *Arabidopsis*. Proteomic data revealed an enrichment of splicing factors among persulfidated proteins and PRMT5 targets. We focused on LSm4, a U6 snRNP component, and GRP7, a splicing modulator. We show that both proteins are persulfidated *in vivo*, independently of PRMT5-targeted arginine residues, whereas H_2_S availability positively influenced their PRMT5-dependent methylation. These findings reveal a connection between H_2_S signaling and PRMT5-dependent methylation, prompting us to explore their convergence in alternative splicing regulation. Transcriptome-wide analyses revealed substantial overlap between genes exhibiting H_2_S-responsive alternative splicing and those with LSm4 methylation-dependent splicing changes. These transcripts were enriched in functions related to RNA processing, immune responses, and stomatal movement. The *prmt5* mutant and plants expressing methylation-deficient LSm4 were impaired in H_2_S-induced stomatal closure. Furthermore, H_2_S-responsive splicing was more pronounced in guard-cell-enriched epidermal samples than in mesophyll-enriched samples. Together, our findings identify PRMT5-dependent LSm4 methylation as a link between H_2_S signaling and alternative splicing, with functional implications for stomatal regulation.

**Highlights:** A gaseous signal, hydrogen sulfide, tunes how plants process RNA by boosting PRMT5-mediated methylation of splicing factors, which contributes to stomatal closure regulation.

## Introduction

Hydrogen sulfide (H_2_S) has emerged as an important redox-reactive signaling molecule in plants, with regulatory functions spanning development, responses to environmental challenges, and plant immunity (Corpas *et al*., 2026). Endogenous H_2_S is produced through multiple enzymatic pathways operating in different subcellular compartments, including the cytosol, chloroplasts, and mitochondria (Aroca *et al*., 2020). In *Arabidopsis thaliana*, L-CYSTEINE DESULFHYDRASE 1 (DES1) represents a major cytosolic source of H_2_S and is induced in response to both abiotic stresses and pathogen infection (Shi *et al*., 2015; Yang *et al*., 2023).

Protein persulfidation is an important mechanism through which H_2_S exerts its signaling functions. This reversible redox post-translational modification (PTM) involves the conversion of protein cysteine thiols (-SH) into persulfides (-SSH), a process that can involve prior oxidation of the thiol group (Vignane and Filipovic, 2023). Persulfidation can affect protein activity, stability, subcellular localization, and molecular interactions, thereby modulating a broad range of cellular processes. Moreover, persulfidation may protect cysteine residues against further oxidation and irreversible oxidative damage (Aroca *et al*., 2021). Plant proteomic studies have identified numerous persulfidated proteins, including transcription factors and RNA-binding proteins (RBPs), suggesting that H_2_S signaling may influence gene expression at multiple levels (Aroca *et al*., 2017; Corpas *et al*., 2025).

Among the mechanisms contributing to gene expression regulation, alternative splicing (AS) plays an important role in expanding transcript diversity and adjusting gene expression in response to developmental and environmental cues (Syed *et al*., 2012; Seok *et al*., 2026). The activity and function of splicing regulators are themselves subject to post-translational control. Indeed, several PTMs, including phosphorylation, methylation, and ubiquitination, modulate splicing factors and ribonucleoprotein complexes, thereby influencing pre-mRNA processing (Deng *et al*., 2010; Lan *et al*., 2022; Rodriguez Gallo and Uhrig, 2023; Agrofoglio *et al*., 2024). In plants, protein arginine methyltransferase 5 (PRMT5)-catalysed symmetric arginine dimethylation is an established mechanism of post-translational regulation targeting several splicing-related proteins, including LSm4, a component of the U6 snRNP, and contributing to the regulation of snRNP function and AS under stress conditions (Deng *et al*., 2010; Sanchez *et al*., 2010; Agrofoglio *et al*., 2024). However, whether H_2_S contributes to AS regulation through interactions with other PTM-dependent mechanisms remains largely unexplored. Here, we investigated whether H_2_S signaling intersects with PRMT5-dependent arginine methylation of splicing-related proteins in the regulation of AS in *A. thaliana*.

## Materials and methods

### Plant material and growth conditions

*A. thaliana* (L.) Heynh. ecotype Columbia-0 (Col-0) was used as the wild type. The *lsm4-1* (SALK_063398; Perez-Santángelo *et al*., 2014) and *prmt5-5*(Sanchez *et al*., 2010)mutant lines are in the Col-0 background. The *des1-1* mutant (SALK_103855), which is defective in the cytosolic H_2_S-producing enzyme L-cysteine desulfhydrase DES1, was used to investigate the role of endogenous H_2_S metabolism (Aroca *et al*., 2017).

To investigate the role of LSm4 arginine methylation, we used *lsm4-1* plants complemented with either wild-type LSm4 or a methylation-deficient variant (LSm4-RxK), in which arginine residues within the nine RGG domains were substituted with lysine. Both untagged and N-terminal YFP-tagged versions of these constructs, expressed under the control of the CaMV 35S promoter, were used (Agrofoglio *et al*., 2024). For GRP7 analyses, we used transgenic lines expressing GFP-tagged wild-type GRP7 or the GRP7-R141K variant in the *grp7-1* background. The GRP7-R141K carries an arginine-to-lysine substitution at residue 141, a PRMT5-targeted methylation site (Steffen *et al*., 2024).

Seeds were surface-sterilized, sown on Murashige and Skoog (MS) medium supplemented with 0.8% (w/v) agar, and stratified in darkness at 4 °C for 3 days. Seedlings were subsequently grown in a controlled-environment growth chamber at 22 °C under long-day conditions (16 hours light/8 hours dark), with a white-light intensity of 80 μmol photons m^−2^ s^−1^ and used at 14 days after sowing. For infection assays involving leaf infiltration, plants were grown in soil at 22 °C under long-day conditions (16 hours light/8 hours dark) with a white-light intensity of 80 μmol photons m^−2^ s^−1^.

### H_2_S treatment of Arabidopsis seedlings

Fourteen-day-old Arabidopsis seedlings grown on MS medium were treated with 150 μM sodium hydrosulphide (NaHS; Sigma-Aldrich, St. Louis, MO, USA), used as an H_2_S donor, or 100 μM hypotaurine (HT; Sigma-Aldrich), used as an H_2_S scavenger. Water was used as a control. Treatments were applied by spraying the Petri dishes containing the seedlings with approximately 1 mL of the corresponding solution (20–25 sprays). Seedlings were incubated for 1 or 4 hours at 22 °C and subsequently harvested, immediately frozen in liquid nitrogen, and stored at −80 °C until further analysis.

### Persulfidation assay

Proteins were isolated from 100 mg of 15-day-old seedlings grown under a long-day photoperiod using extraction buffer (PBS 1X, pH 7.4, 1 mM EDTA, 2% SDS, 1 mM phenylmethylsulfonyl fluoride (PMSF), and protease inhibitor cocktail (Sigma-Aldrich, USA). For *in vitro* persulfidation assays, protein extracts were incubated with freshly prepared 200 μM NaHS, either in the presence or absence of 10 mM DTT (Merck), for 10 minutes at room temperature in the dark. For in vivo persulfidation assays, no NaHS was added to the protein extracts. Subsequently, persulfidation was assessed using the dimedone switch method as described by Aroca *et al*., 2022. Following labeling with DCP-bio (Xoder Technologies), the protein pellet was resuspended in immunoprecipitation buffer containing 50 mM Tris-HCl, pH 7.2, 100 mM NaCl, 10% glycerol, 0.1% Tween™ 20, 1 mM PMSF, and protease inhibitor cocktail (Sigma-Aldrich, USA). Protein extracts were incubated with pre-washed NeutrAvidin-Magnetic beads (Thermo Fisher Scientific) at 4°C for 1 hour with gentle rotation. Input samples collected before purification were saved as controls. Beads were washed four times with IP buffer, and bound proteins were eluted in reducing SDS-PAGE sample buffer. Persulfidated YFP-LSM4 and YFP-LSM4RxK proteins were identified by immunoblotting with α-GFP, while GRP7 and GRP7R141K proteins were identified using α-GRP7 (Streitner *et al*., 2008).

### Protein extraction and Immunoblot analysis

Total proteins were extracted using an extraction buffer containing 50 mM Tris-HCl (pH 7.2), 100 mM NaCl, 10% (v/v) glycerol, 0.1% (v/v) Tween-20, 1 mM phenylmethylsulfonyl fluoride (PMSF), and a plant protease inhibitor cocktail (Sigma). Extracts were clarified by centrifugation at 12,000 × g for 15 minutes at 4°C. Equal amounts of protein samples were separated by SDS-PAGE and transferred onto nitrocellulose membranes. Membranes were incubated overnight with the following primary antibodies: anti-GFP (1:4,000; Invitrogen, Cat. No. A6454), anti-GRP7 (Streitner *et al*., 2008), SYM10 (1:1.000; Millipore, Cat. No. 07-412). Membranes were then incubated with a peroxidase-conjugated anti-rabbit secondary antibody (1:5,000; Invitrogen, Cat. No. 31460). Immunoreactive proteins were detected using an ECL detection kit (Pierce) and an ImageQuant LAS 4000 system (GE Healthcare). Ponceau S staining of the large Rubisco subunit was used to assess protein loading.

### Immunoprecipitation and methylation analysis

Total protein extracts were prepared from2-week-old *GRP7:GRP7-GFP* seedlings using an immunoprecipitation buffer composed of 50 mM Tris-HCl (pH 7.5), 150 mM NaCl, 10% glycerol, 0.1% Nonidet™ P40, 1 mM phenylmethylsulfonyl fluoride (PMSF), and protease inhibitor cocktail (Sigma-Aldrich, USA). Following centrifugation at 12,000 × g for 15 minutes at 4°C, 500 μg of total protein were incubated with GFP-Trap Agarose beads (Chromotek, Germany) for 1 hour at 4°C under gentle agitation. The beads were subsequently collected by centrifugation at 2,500 × g for 4 minutes, and an aliquot of the supernatant was retained as the flow-through fraction. After four washes with immunoprecipitation buffer, bead-bound proteins were released by resuspension in 2.5× Laemmli SDS sample buffer. GRP7 arginine methylation was assessed by immunoblotting with the anti-SYM10 antibody, as described above.

### RNA extraction and RT-PCR analysis

Total RNA was extracted from two-week-old *Arabidopsis* seedlings and from guard cell-or mesophyll cell-enriched tissues obtained from five-to six-week-old plants, using TRIzol reagent (Ambion) according to the manufacturer’s instructions. RNA samples were treated with RNase-free DNase I (Promega) to remove residual genomic DNA. For reverse transcription, 1 µg of total RNA was converted into cDNA using SuperScript II reverse transcriptase (Invitrogen). The resulting cDNA was diluted 1:3 and used for RT-PCR analysis of alternative splicing events. PCR amplification was performed using Platinum SuperFi III DNA Polymerase-High Fidelity (Invitrogen) and gene-specific primers. The primer sequences used to analyse alternative splicing events in AT1G72900 (AtTN7) were forward (F): 5′-CCGCTTTGGCAGATATCTCC-3′ and reverse (RV): 5′-CCTCCCATTGCTTATTCTTG-3′. For AT1G32230 (RCD1), two separate PCR reactions were performed using a common reverse primer combined with either forward primer F1 or F2. The F1/RV primer pair was used to amplify the intron-retaining isoform, whereas the F2/RV primer pair amplified the spliced isoform. The primer sequences were as follows: F1: 5′-GAAGTCCGATGACATACACTGG-3′; F2: 5′-CCCCAATGCTGAAGGTAATTTGAT-3′; and RV: 5′-CTCTTGCTCCCTGGTTTGACTC-3′. PCR products were separated by electrophoresis on 2.5% (w/v) agarose gels and visualized using SYBR Green.

For intron retention (IR) quantification, non-saturated gel images were acquired in TIFF format. Band intensities were quantified using Fiji/ImageJ. For each sample, a rectangular region of interest (ROI) encompassing the band of interest was selected, and the local background intensity was subtracted. The relative intron retention level was calculated as the ratio of the background-corrected intensity of the intron-retaining band to the sum of the intensities of the intron-retaining and spliced bands: IR/total = I(IR) / [I(IR) + I(spliced)] where I(IR) and I(spliced) represent the background-corrected intensities of the intron-retaining and spliced bands, respectively.

### Proteomic, RNAseq and Gene Ontology (GO) analysis

Information on the publicly available proteomic and RNA-seq datasets, together with the intersections between the datasets shown in Figure 1A and Figure 2A, B, is provided in Supplementary Table S1. GO enrichment was performed on genes displaying differential alternative splicing (DAS) shared between the *prmt5* mutant and both H_2_S RNA-seq datasets, using PANTHER via Fisher’s exact test with FDR correction (Benjamini-Hochberg) and *A. thaliana* as the reference. Significant terms (FDR < 0.05) were processed with REVIGO (SimRel, Dispensability < 0.2) to reduce semantic redundancy, manually prioritizing, among hierarchically related terms, those of greater biological relevance to the phenotype under study. Visualization was generated in R with ggplot2.

**Figure 1.**
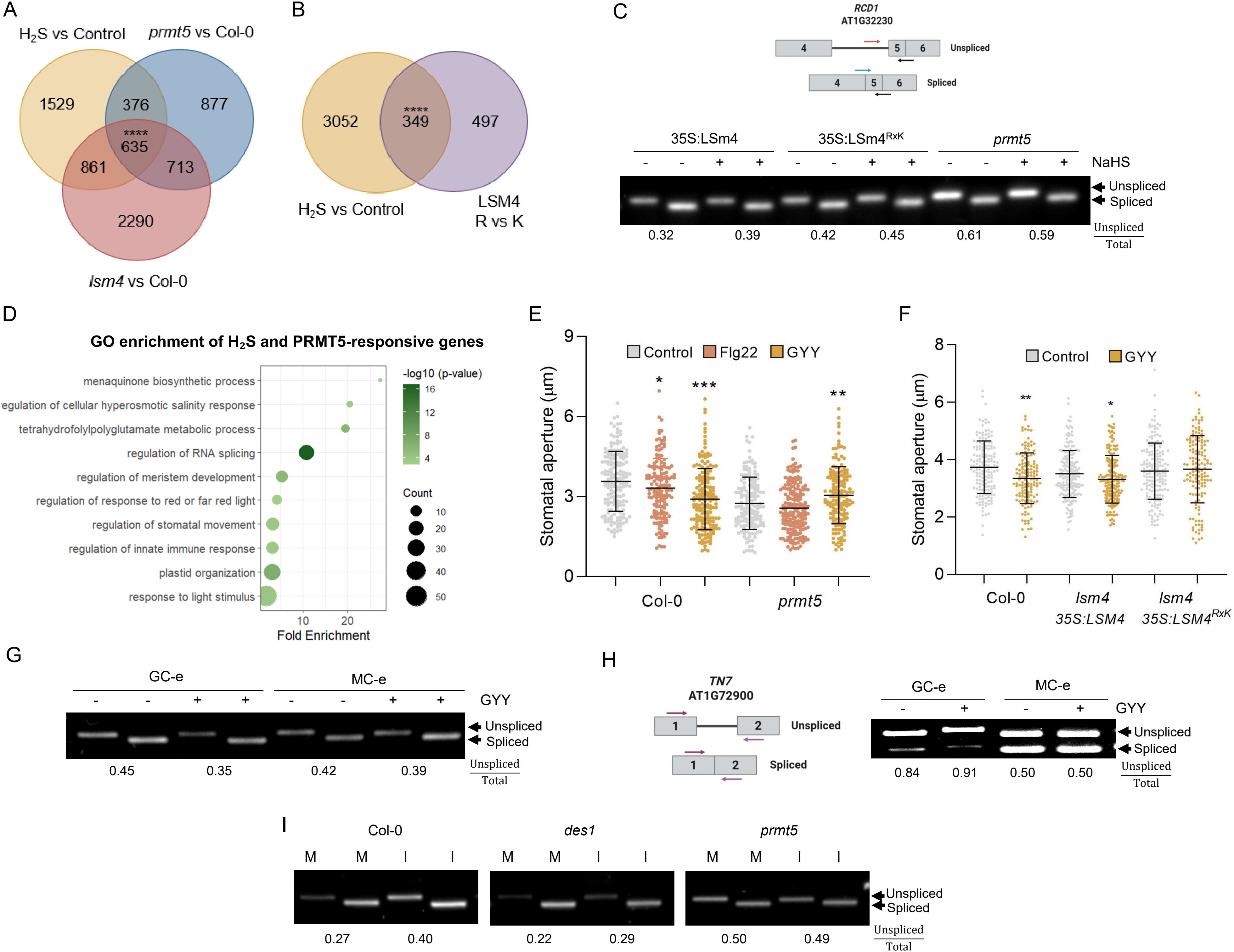
Convergence of persulfidation and PRMT5-dependent methylation on splicing factors. **A-** Upset plot showing theenrichment of splicing factors and PRMT5-targets in SSH-proteomic public datasets ofArabidopsis plants. Enrichment was assessed using a hypergeometric test ***\*\**** *p < 0*.*01; **** p <0*.*0001*.**B–C**. *In vivo* persulfidation of YFP-LSm4, YFP-LSm4^RXK^ (B), GRP7, and GRP7^R141K^ (C), assessed by the dimedone-switch method followed by NeutrAvidin purification and immunoblotting (IB) using anti-GFP (B) or anti-GRP7 (C) antibodies. Input fractions before purification are shown as controls. Representative blots are shown (two biological replicates, each comprising n = 30 plants per line) **D-** Detection of LSm4 arginine symmetric dimethylation in Col-0, *prmt5* and *lsm435S:LSm4-YFP* by immunoblot with the Sym10 antibody. Ponceau red staining of the large subunit of the Rubisco (RbcL) shows loading across samples. The arrows indicate expected size for LSm4 and LSm4-YFP. Meth: symmetrically dimethylated arginine. **E-** LSm4 arginine symmetric dimethylation in Col-0 and *des1* plants in the presence or absence of H_2_S exogenous treatment with ⍰150 M NaHS for 1 hour. Ponceau red staining of the large subunit of the Rubisco (RbcL) shows equal loading across samples. **F-** Quantification of (E). Two-way ANOVA followed by Tukey’s multiple comparisons test. Different letters indicate significant differences (*p*< 0.05; five biological replicates, each comprising n = 30 plants per line and treatment). **G-** LSm4 arginine symmetric dimethylation in Col-0 plants in the presence or absence of H_2_S scavenger, hypotaurine 100 M for 1 hour. Ponceau red staining of the large subunit of the Rubisco (RbcL) shows equal loading across samples. **H-** Quantification of (G). t-test, * *p*<0.05. (three biological replicates, each comprising n = 30 plants per line). **I-** GRP7-GFP arginine symmetric dimethylation in the presence or absence of ⍰150 M NaHS for 1 hour. GRP7-GFP was immunoprecipitated (IP) with GFP-trap, GRP7-GFP methylation and input levels were detected by IB with Sym10 (top) or⍰-GFP antibody (bottom), respectively. Representative blots are shown (two biological replicates, each comprising n = 30 plants per treatment).

**Figure 2.**
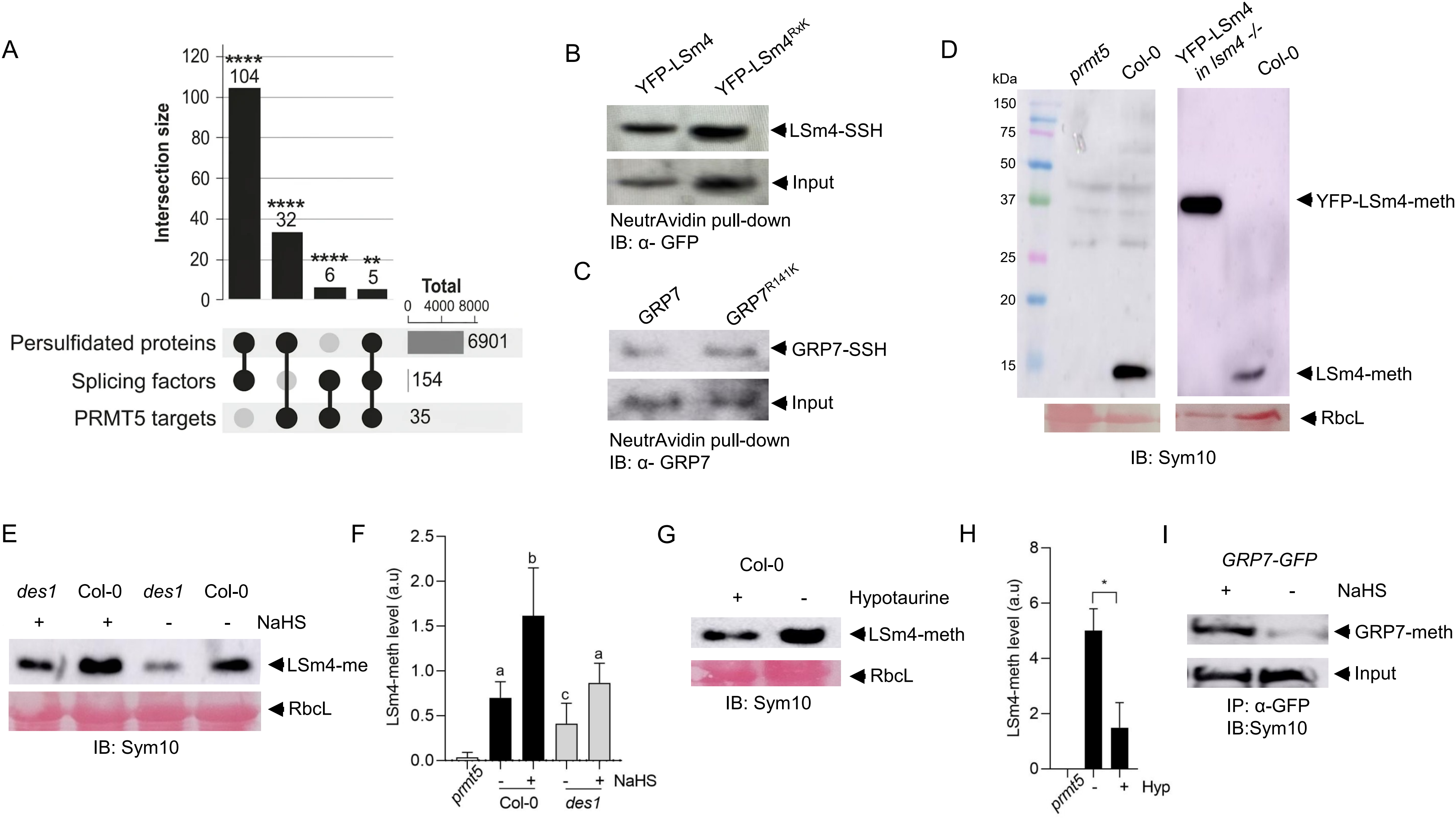
H_2_S–PRMT5–LSm4 signaling regulates alternative splicing and stomatal responses during plant immunity. **A–B**. Venn diagrams showing the overlap between genes with pre-mRNA splicing events altered in *lsm4* and *prmt5* mutants and those altered following exogenous treatment with the H_2_S donors (A), and between genes with splicing events affected by LSm4 methylation status and those altered by NaHS treatment (B). Statistical significance of the overlaps was assessed using a hypergeometric test. *\*\*\*\* p*< 0.0001. **C-** RT-PCR analysis of intron 4 retention in *RCD1* in *lsm4 35S:LSM4 and lsm4 35S:LSM4*^*RxK*^ transgenic lines and *prmt5* mutants treated with 150 μM NaHS for 4 hours. The alternative splicing event is schematized above the gel, with primer positions indicated. The ratio of intron-retaining transcript to total transcript is shown. **D**. Gene Ontology (GO) enrichment analysis of genes showing overlapping splicing alterations in *prmt5* mutants and following H_2_S treatment, represented as a bubble plot. Gene count in each category is represented by the size of the circles, and color intensity represents the p-value. **E-** Stomatal aperture measurements in epidermal strips from Arabidopsis Col-0 and *prmt5* mutants treated with 100 nM flg22 or 100 μM GYY4137 for 90 min, after preincubation for 3 hours in opening buffer (5 mM MES pH 6.1, 50 mM KCl) under light. **F-** Stomatal aperture measurements in epidermal strips from Arabidopsis Col-0, *lsm4 35S:LSM4 and lsm4 35S:LSM4^RxK^* lines. Epidermal strips were preincubated for 3 hours in the opening buffer under light and subsequently treated with 100 μM GYY4137 for 90 min under light. **(E, F)** The values of stomatal aperture are expressed in microns (µm). (Four independent experiments, n= approximately 50 stomata per genotype per treatment). Statistical significance was determined by two-way ANOVA followed by Dunnett’s post hoc multiple comparisons against the control. *\*p*< 0.05, *\*\*p*< 0.01, *\*\*\*p*< 0.001. **G-** RT-PCR analysis of intron 4 retention in *RCD1* in guard cell-enriched (GC-e) and mesophyll-enriched (MC-e) RNA samples from wild-type plants. Epidermal strips and mesophyll tissue were preincubated for 3 hours in opening buffer (10 mM K-MES, pH 6.1, and 10 mM KCl) and subsequently treated with or without 100 μM GYY4137 (H_2_S donor) for 90 min. **H-** RT-PCR analysis of the alternative splicing event in *TN7* in guard cell-enriched (GC-e) and mesophyll-enriched (MC-e) RNA samples from wild-type plants treated with or without 100 μM GYY4137 for 90 min, after preincubation in opening buffer as described in (G). **I-** RT-PCR analysis of the alternative splicing event in *RCD1* in Col-0, *des1* and *prmt5* mutants 24 hours after *Pseudomonas syringae* DC3000 (I) or mock treatment (M).

### Infection assay

Two-week-old Arabidopsis plants were inoculated with a bacterial suspension *of Pseudomonas syringae* pv. *tomato* DC3000 (OD_600_ = 0.002) in 10 mM MgSO_4_ according to Agrofoglio *et al*., 2026. Mock plants were treated with 10 mM MgSO_4_. Leaves were harvested at 1-day post-inoculation (dpi).

### Guard cell analysis

#### Stomatal aperture assay

Stomatal aperture assays were performed according to Pantaleno *et al*., 2024. Briefly, epidermal peels from abaxial side of fully expanded leaves from five to six-week-old Arabidopsis plants were excised with tweezers and immediately floated in opening buffer (5 mM MES pH 6.1, 50 mM KCl) for 3 hours under light (200 μmol photons m^−2^ s^−1^) and subsequently maintained in the same buffer (Control) or exposed to different treatments as indicated in the legends. Stomata were photographed using an AmScope MU1000 camera coupled to an Olympus CKX53 microscope with a 40× lens (LUCPlanFLN, 0.6 numerical aperture). The stomatal aperture width was quantified using the ImageJ analysis software (NIH, Bethesda, MD, USA).

#### Guard cell- and mesophyll-enriched tissue collection

Guard cell–enriched (GC-e) and mesophyll cell–enriched (MC-e) tissues were obtained according to Pantaleno *et al*., 2024. Briefly, for GC-e samples, epidermal peels from the abaxial side of fully expanded Arabidopsis leaves were collected with tweezers and placed in an opening buffer-containing Petri dish until the surface of the dish is fully covered with peels. The peels were incubated for 3 hours in white light and then the pharmacological treatments were applied. For MC-e samples the rest of the leaf without abaxial epidermal peel was incubated in the opening buffer in a Petri dish as GC-e samples. A total of 12-15 plants per treatment, per replicate were used.

### Statistical analysis

All experiments were performed using independent biological replicates, as indicated in the corresponding figure legends. Technical replicates were treated as subsamples and were not considered independent biological replicates. Data are presented as mean ± standard error (SE). Statistical analysis was conducted as described in the figure legend using Graphpad Prism version 5.01 software.

## Results and Discussion

### Protein persulfidation is enriched in splicing factors regulated by PRMT5

We first analyzed publicly available *A. thaliana* datasets of protein persulfidation to explore a potential link between H_2_S signaling and post-transcriptional gene regulation (Supplemental Table S1). This analysis revealed a substantial overlap between persulfidated proteins and RNA-binding proteins involved in pre-mRNA processing, including spliceosomal components and auxiliary splicing modulators which are commonly referred to as splicing factors (SF; Fig. 1A). Among the 154 curated SFs analyzed(Reddy *et al*., 2013), 104 were identified as persulfidated in at least one large-scale proteomic dataset, accounting for approximately 70% of this group. Consistently, persulfidated proteins were significantly enriched among SFs compared with their representation in the Arabidopsis proteome. In line with this enrichment, targeted studies have provided evidence of persulfidation in specific SFs, including U2 snRNP auxiliary factor 65A (AtU2AF65a) (Ma *et al*., 2024, 2025)and RNA-binding motif protein 25 (AtRBM25) (Wu, *et al*., 2026, b) in *A. thaliana*. In addition, persulfidation was demonstrated for alternative splicing factor ATO (BraATO2) in *B. rapa* and detected *in vitro* for its Arabidopsis homolog, AtATO (Hao *et al*., 2025). Notably, SFs were also enriched among proteins undergoing nitric oxide-dependent PTMs across diverse eukaryotic systems, including plants, invertebrates, fish, and mammals (Schindler *et al*., 2026), suggesting that redox-dependent PTMs broadly target RNA-processing factors across eukaryotes.

Recent evidence indicates that PTMs of SFs can alter their regulatory activity and reshape alternative splicing (Kretova *et al*., 2023; Agrofoglio *et al*., 2024). Moreover, the convergence of different PTMs on RNA-processing factors may create regulatory nodes linking cellular signaling to changes in RNA processing. Consistent with this emerging paradigm, a subset of the persulfidated splicing factors identified in *A. thaliana* had been previously reported as targets of symmetric arginine dimethylation catalyzed by protein arginine methyltransferase 5 (PRMT5) (Deng *et al*., 2010; Z. Zhang *et al*., 2011; P. Hu *et al*., 2019; Cao *et al*., 2022; Martín-Merchán *et al*., 2024). Although PRMT5 targets comprise functionally diverse proteins, 32 of the 35 previously reported PRMT5 targets were also identified as persulfidated proteins (Fig. 1A). This overlap was particularly pronounced among SFs (Fig. 1A). All six splicing-related components previously reported as PRMT5 targets were detected in the persulfidated proteomes (Fig. 1A), raising the possibility that individual RNA-processing factors may be subject to regulation by both cysteine persulfidation and arginine methylation. Given the established role of PRMT5 in snRNP biogenesis, spliceosome activation, and alternative splicing regulation (Deng *et al*., 2010, 2016; Sanchez *et al*., 2010), these observations provided a rationale to investigate whether persulfidation and arginine methylation converge to modulate RNA processing and alternative splicing in plants.

### H_2_S signaling regulates PRMT5-dependent methylation of LSm4 and GRP7

Among the proteins identified in both the persulfidation and PRMT5-dependent methylation datasets (Fig. 1A), we focused on LSm4, a core component of the U6 snRNP spliceosome complex, and GRP7, an RNA-binding protein also shown to have a limited role in splicing. Both are well-characterized targets of PRMT5-mediated symmetric arginine dimethylation (Deng *et al*., 2010), and we previously showed that their methylation is relevant for plant adaptation to stress (Agrofoglio *et al*., 2024; Steffen *et al*., 2024). Given that LSm4 and GRP7 were identified as persulfidated targets in large-scale datasets obtained under high H_2_S donor concentrations (Romero *et al*., 2026), we first asked whether both proteins also undergo persulfidation under physiological conditions, and whether this modification depends on their methylation status. We evaluated persulfidation status using a dimedone-switch assay (Aroca *et al*., 2022). Persulfidation of LSm4 and GRP7 was readily detected *in vivo* (Fig. 1B and C), indicating that these proteins undergo persulfidation by endogenous intracellular H_2_S levels. As a control for the specificity of the biotin signal detected after dimedone-switch assay, persulfidation signals were reversed by DTT (Supplemental Figure S1). Next, to assess whether persulfidation depends on PRMT5-mediated methylation, we compared wild-type LSm4 or GRP7 with variants in which we previously demonstrated that the PRMT5-targeted arginine residues were substituted by lysines, thereby preventing their methylation (Agrofoglio *et al*., 2024; Steffen *et al*., 2024). Both proteins displayed comparable levels of persulfidation relative to the input irrespective of their methylation status (Fig. 1B and C), indicating that persulfidation of LSm4 and GRP7 does appear to require prior methylation at the PRMT5-targeted arginine residues.

In the opposite direction, we examined whether H_2_S signaling influences LSm4 methylation. To this end, LSm4 methylation was assessed by immunoblotting using the Sym10 antibody, which recognizes PRMT5-dependent symmetric arginine dimethylation. A methylation signal corresponding to endogenous LSm4 was readily detected in wild-type seedlings but was absent in *prmt5* mutants. Consistent with the identity of this signal, analysis of *lsm4* plants complemented with YFP-LSm4 revealed a shift toward a higher-molecular-weight band corresponding to the fusion protein (Fig. 1D). We next investigated the effect of H_2_S on LSm4 methylation using complementary genetic and pharmacological approaches. In the *des1* mutant, which is impaired in one of the major sources of endogenous cytosolic H_2_S production, LSm4 methylation was reduced, whereas treatment of Col-0 plants with the H_2_S donor NaHS increased LSm4 methylation (Fig. 1E–F). Importantly, NaHS treatment restored LSm4 methylation in *des1* plants to levels comparable to those observed in untreated Col-0 plants (Fig. 1E-F). Consistent with these findings, depletion of endogenous H_2_S using hypotaurine, an H_2_S scavenger, also reduced LSm4 methylation (Fig. 1G, H). To determine whether the positive effect of H_2_S on protein methylation also extended to GRP7, we immunoprecipitated GRP7-GFP from 35S plants treated with or without the H_2_S donor NaHS. NaHS treatment increased GRP7 methylation levels (Fig. 1I), indicating that H_2_S-mediated enhancement of protein methylation is not restricted to LSm4. Together, this genetic and pharmacological evidence indicates that H_2_S availability positively regulates PRMT5-dependent methylation of LSm4 and GRP7. Reciprocal regulation is also possible. L-cysteine desulfhydrase (LCD), another enzyme involved in H_2_S production, has been reported as a PRMT5 substrate *in vitro*, with its methylation enhancing enzymatic activity under cadmium stress (Cao *et al*., 2022). These findings raise the possibility of a reciprocal interplay between H_2_S signaling and PRMT5-dependent methylation. On one hand, H_2_S may influence PRMT5-dependent substrate methylation through redox-dependent regulation of PRMT5 itself and/or its substrates. In this regard, PRMT5 activation by another redox signal, nitric oxide (NO), has also been reported (Hu *et al*., 2017). On the other hand, PRMT5-mediated methylation of LCD may enhance its enzymatic activity and thereby promote H_2_S production, potentially establishing a feedback mechanism linking redox signaling and protein arginine methylation. Interestingly, a connection between methyl metabolism and H_2_S biosynthesis has also been described in mammals. S-adenosylmethionine (SAM), the methyl donor used by protein arginine methyltransferases, acts as an allosteric activator of human cystathionine β-synthase (CBS), an enzyme involved in H_2_S production (Ereño-Orbea *et al*., 2014). Although mechanistically distinct from the regulatory interplay proposed here, this example further illustrates the connection between methyl metabolism and H_2_S biosynthesis.

### H_2_S-responsive alternative splicing converges with the PRMT5–LSm4 pathway

Having established that H_2_S positively regulates PRMT5-dependent methylation of LSm4, we next asked whether this molecular crosstalk translates into changes in alternative splicing at the transcriptome level. To address this question, we first compared genes displaying altered splicing in *prmt5* and *lsm4* mutants (Mateos *et al*., 2023; Agrofoglio *et al*., 2024)with those affected by exogenous H_2_S treatment in two independent datasets (Ma *et al*., 2024; Wu *et al*., 2026, b). A substantial overlap was observed among the three datasets, with 635 genes showing altered splicing in all three conditions (Fig. 2A). Notably, the *prmt5* mutant displayed altered splicing in 2,601 genes, of which approximately 39% (1,011/2,601) were also differentially spliced following H_2_S treatment. Of these, 63% (635/1011) also showed altered splicing in the *lsm4* mutant, further connecting H_2_S- and PRMT5-dependent splicing programs with LSm4 function. We next investigated whether this convergence was specifically associated with transcripts whose splicing depended on LSm4 methylation (Agrofoglio *et al*., 2024). Among these 846 genes, 349 (41%) were also H_2_S-responsive (Fig. 2B). Together, these results support an association between H_2_S signaling, PRMT5 activity, and LSm4 methylation in the regulation of alternative splicing.

To experimentally validate the convergence identified through the comparative analysis of independent splicing datasets, we selected an intron retention (IR) event in *RADICAL-INDUCED CELL DEATH1* (*RCD1*), a plant-specific regulator implicated in the control of defense-related responses (Zhu *et al*., 2013; Wirthmueller *et al*., 2018; Xing *et al*., 2026). We analyzed *RCD1* splicing following exogenous H_2_S treatment in plants expressing wild-type LSm4, the methylation-deficient LSm4RxK variant, and *prmt5* mutants. H_2_S treatment increased intron retention in *RCD1* in plants expressing wild-type LSm4, whereas this response was not observed in plants expressing LSm4RxK or in *prmt5* mutants (Fig. 2C). These results provide experimental support for the involvement of PRMT5 activity and LSm4 methylation in H_2_S-responsive *RCD1* splicing.

### PRMT5-mediated LSm4 methylation contributes to H_2_S-dependent stomatal closure

To explore the biological processes associated with the convergence of H_2_S- and PRMT5-dependent splicing regulation, we performed a Gene Ontology (GO) enrichment analysis of genes displaying altered splicing in both conditions. GO analysis revealed enrichment in the category “regulation of RNA splicing” (Fig. 2D), consistent with the broad effects that perturbing splicing regulators has on RNA-processing networks (Agrofoglio *et al*., 2024, 2026). Notably, 68% of the persulfidated splicing factors identified in Fig. 1A were themselves H_2_S-responsive at the splicing level (Supplemental Table S1). This overlap suggests that H_2_S may act on splicing factors at two interconnected levels, through persulfidation of the proteins and through regulation of their own pre-mRNA splicing, which could give rise to feedback loops. Such a scenario is consistent with evidence that, in plants, alternative splicing of a splicing factor can generate non-coding isoforms that modulate the splicing of the factor itself in response to environmental cues (Rodríguez *et al*., 2026, Preprint).

Beyond RNA-processing categories, “regulation of stomatal movement” and “regulation of innate immune response” were among the enriched biological processes (Fig. 2D). In addition, the term “regulation of cellular response to hyperosmotic salinity” was enriched almost exclusively through the chloroplast K^+^/H^+^ antiporters KEA1, KEA2 and KEA3, implicated in ROS metabolism and stomatal responses (Sánchez-McSweeney *et al*., 2021; Yamada *et al*., 2024). These enrichments prompted us to investigate whether the H_2_S–PRMT5–LSm4 pathway contributes to the regulation of stomatal responses, particularly during immune signaling. To this end, we examined stomatal aperture following treatment with either an H_2_S donor or the bacterial elicitor flg22, which induces H_2_S-dependent stomatal closure (Scuffi *et al*., 2025). As expected, Col-0 plants efficiently closed their stomata in response to both treatments, whereas *prmt5* mutants failed to close their stomata in response to flg22 and, remarkably, exhibited increased stomatal aperture rather than closure following exogenous H_2_S treatment (Fig. 2E). Interestingly, a similar reversal of the stomatal response has been reported in *des1* mutants, which exhibit increased stomatal aperture rather than closure in response to high flg22 concentrations(Scuffi *et al*., 2025). Importantly, the *prmt5* mutant retained the ability to close their stomata in response to a NO donor, indicating that their altered responses to H_2_S and flg22 do not reflect a general impairment of stomatal movement (Supplemental Figure S2A). Then, in order to assess whether PRMT5-dependent methylation of LSm4 contributes to H_2_S-induced stomatal closure, we compared plants expressing wild-type LSm4 with those expressing the methylation-deficient LSm4RxK variant in a *lsm4* background. Plants expressing LSm4 showed stomatal closure in response to the H_2_S donor, resembling the wild-type response (Figure 2F). In contrast, plants expressing LSm4RxK failed to close their stomata under the same conditions (Figure 2F). These results indicate that the PRMT5-targeted arginine residues of LSm4 are required for a normal stomatal response to exogenous H_2_S and support a role for LSm4 methylation in H_2_S-dependent stomatal closure.

### H_2_S–PRMT5–LSm4 signaling links alternative splicing and stomatal responses during plant immunity

Recent studies have shown that H_2_S-mediated regulation of alternative splicing can generate isoforms with differential effects on stomatal dynamics in Arabidopsis and *B. rapa* under ABA treatment and drought-related stress (J. Zhang *et al*., 2025; Wu *et al*., 2026, *a*). In addition, alternative splicing is increasingly recognized as an important regulatory layer in plant immunity, where the abundance of immune receptors must be tightly regulated to allow their induction upon pathogen recognition while preventing inappropriate activation under non-infected conditions (Iglesias *et al*., 2026). These findings raise the possibility that H_2_S-dependent alternative splicing contributes to the regulation of stomatal responses and plant immunity.

To explore this possibility, we examined whether H_2_S regulates alternative splicing in guard cells, which play a central role in both stomatal regulation and plant defense. To this end, RNA was isolated from epidermal strips which provide a guard cell-enriched preparation (Pantaleno *et al*., 2024). The relative expression levels of *ECERIFERUM2* (*CER2*; Leonhardt *et al*., 1999; Wang *et al*., 2011)and *RBCS2B* (*AT5G38420*; Sawchuk *et al*., 2008), used as markers for guard cells and mesophyll cells, respectively, confirmed the enrichment of the corresponding cell populations (Supplemental Figure S2B). We then evaluated splicing isoforms of *RCD1* and *TN7*, which encodes a TIR-NBS-LRR (TNL) immune receptor whose splicing is regulated by *Pseudomonas* infection and LSm4 methylation (Agrofoglio *et al*., 2024). Interestingly, H_2_S donor treatment had a more pronounced effect on *RCD1* splicing in guard-cell-enriched samples than in mesophyll-enriched samples (Figure 2G). For *TN7*, intron retention (IR) was more prominent in guard-cell-enriched samples than in mesophyll-enriched samples under control conditions, suggesting cell-type-specific differences in its basal splicing pattern. Furthermore, H_2_S donor treatment altered *TN7* splicing in guard-cell-enriched samples but had no detectable effect in mesophyll-enriched samples (Figure 2H).

Finally, to explore whether H_2_S biosynthesis and PRMT5 function contribute to the pathogen-induced splicing regulation of *RCD1*, intron retention was analyzed in Col-0, *des1*, and *prmt5* plants 24 hours after inoculation with *P. syringae* DC3000 or mock treatment (Figure 2I). In Col-0 plants, bacterial treatment increased IR in *RCD1* relative to mock-treated controls. In contrast, neither *des1* nor *prmt5* mutants showed changes in IR by *P. syringae* exposure. Despite distinct basal splicing profiles, neither mutant recapitulated the infection-induced splicing changes observed in wild-type plants.

Collectively, our findings reveal a previously unrecognized connection between H_2_S signaling, PRMT5-dependent protein methylation, and alternative splicing. The extensive overlap among H_2_S-, PRMT5-, and LSm4-dependent splicing programs, together with our physiological analyses, supports a model in which H_2_S influences alternative splicing, at least in part through the regulation of LSm4 methylation, with potential implications for stomatal responses during plant immunity. Our work thus points to a potential molecular link between redox signaling and splicing regulation, suggesting that alternative splicing may represent one of the mechanisms through which H_2_S influences plant immune responses.

## Abbreviations

(AS): Alternative splicing
(GRP7): Glycine-rich RNA-binding protein 7
(GC-e): Guard cell–enriched
(H_2_S): Hydrogen sulfide
(DES1): L-cysteine desulfhydrase 1
(MC-e): Mesophyll cell–enriched
(PTM): Post-translational modification
(PRMT5): Protein arginine methyltransferase 5
(RBPs): RNA-binding proteins
(LSm4): Sm-like protein 4
(SF): Splicing factor

## Supplementary data

**Supplemental Figure S1:** DTT reverses persulfidation signals detected by the dimedone-switch assay. (A) total protein (B) GRP7

**Supplemental Figure S2:** Validation of tissue enrichment and NO-induced stomatal closure in prmt5 mutants. A) Expression of guard cell and mesophyll cell marker genes (*ECERIFERUM2*, CER*2*, and *RBCS2B*, respectively) in guard-cell-enriched and mesophyll-enriched samples. (B) Stomatal aperture in Col-0 and *prmt5* mutants following treatment with the NO donor, SNP 100 M.

**Supplemental Table S1**. Publicly available proteomic and RNA-seq datasets used to identify protein and transcript overlaps related to H_2_S signaling, PRMT5, and alternative splicing in Arabidopsis thaliana. (A) Publicly available proteomic and transcriptomic datasets of PRMT5 substrates, splicing factors and accessory splicing regulators, H_2_S-responsive splicing factors, and persulfidated proteins, including their overlapping members. (B) Publicly available RNA-seq datasets used to identify genes showing alternative splicing in *lsm4* and *prmt5* mutants, LSm4 RxK-expressing plants in the *lsm4* background, and H_2_S-treated plants, including gene-level overlaps among datasets. (C) Public Dataset References.

## Acknowledgements

SBT, YCA, DS, CGM, RP, JLM, and MJI were supported by the Consejo Nacional de Investigaciones Científicas y Técnicas (CONICET), Argentina. BDL was supported by the Agencia Nacional de Promoción de la Investigación, el Desarrollo Tecnológico y la Innovación (ANPCyT), Argentina. We thank the laboratory of Prof. Dr. Dorothee Staiger (Universität Bielefeld, Germany) for kindly providing the GRP7 transgenic lines and the anti-GRP7 antibody. This work was conducted in the context of severe and sustained underfunding of the Argentine scientific system, which has significantly affected research institutions, scientific infrastructure, and the conditions required to sustain scientific activity

## Author contributions

SBT, DS, RP, CGM, YCA and MBL performed experiments. SBT, DS, and MJI analysed data. SBT and MJI contributed to writing the original draft. YCA, DS, JLM and CGM contributed to manuscript review and editing. JLM and MJI conceived and designed the research. All authors read and approved the final manuscript.

## Data availability

All data supporting the findings of this study are available within the article and its supplementary materials.

